# sMAP: An interactive microarray data analysis tool for early-stage researchers

**DOI:** 10.1101/2022.05.27.492984

**Authors:** Samuel Bharti, Nikita Krishnan, Arian Veyssi, Maryam Momeni, Sneha Raj

## Abstract

Microarray data enables biologists to extract differentially expressed genes (DEGs) across multiple phenotypes. While several pipelines and tools exist to perform microarray data analysis, they are targeted to users with moderate to advanced computational understanding and lack an easy-to-use, interactive and dynamic methodology to perform analysis assisted with comprehensive learning resources. In this study, we developed an interactive application “sMAP” (Standard Microarray Analysis Pipeline) to make transcriptome microarray data analysis more accessible in learning environments and to enable the identification of significant pathological biomarkers. In a case study of colorectal cancer, we showed that sMAP enabled us to reproduce previous findings and discover relevant pathways. sMAP provides a comprehensive set of tutorials and learning documentation to help early-stage researchers. The latest URLs of sMAP’s hosting, tutorial, and frequently updated documentation can be found at https://github.com/BI-STEM-Away/sMAP.

## Introduction

Microarray analysis is a widely used computational technique to identify differentially expressed genes and perform downstream analysis. DNA microarrays utilize glass microscope slides that have many designated indents which contain DNA probes that correspond to genes of interest. mRNA transcripts are extracted from tissues and flow through the microarrays, hybridizing with corresponding DNA probes [1, 2, 3]. This technology has enabled massive data collection that has revolutionized biological experimentation, and the advent of bioinformatics technology has served to make analysis much more efficient and effective. For instance, reliance on DNA sequencing at a large scale is an extra step no longer necessary with microarray analysis [4].

Microarray data analysis provides a challenge to current biological researchers due to a lack of training on the analysis process and the high cost of the software. However, the ability to derive biological inferences from microarray data with thousands of gene expression data points has proven to be useful [5]. Making sense of genomic data allows researchers to identify key disease pathways and find potential therapeutic targets. For example, gene expression profiling has recently uncovered associations between alcohol and neurodegenerative diseases [6], regulon expression in cognitive aging [7], and chamber-specific transcriptomics in heart failure patients [8].

Few tools exist that primarily focus on microarray data analysis, which allows for full sequencing of the transcriptome [9]. sMAP is an R Shiny application for microarray analysis that identifies key steps within the transcriptomics pipeline: quality control and preprocessing, statistical analysis, and functional analysis. sMAP targets early-stage researchers by providing an easy-to-use interface where users can easily navigate through the analysis pathway and visualize the results of statistical and functional analysis.

Current tools, such as iGEAK, MetaOmGraph, and UTAP, have often treated microarray data as a series of separate steps that focus only on data visualization and statistical analysis. Therefore, these tools fail to provide biomedical researchers with the ability to discern the significance of steps in the pipeline and to make biological inferences from their data. MetaOmGraph and UTAP require a prior understanding of R programming so that users can specify gene samples or specifications to parameters for analysis which could be a limitation to early-stage researchers [10, 11]. While tools like iGEAK attempt to address the need for an advanced understanding of R by creating an easily-installed desktop application, sMAP addresses the same limitation of software knowledge by opting for an AWS-hosted web application [12]. Web-based applications are more advantageous to early-stage researchers need tools that are flexible on local and remote computers.

## Methods

sMAP contains four major analysis steps, i.e., Data Importation, Quality Control & Preprocessing, Sample Grouping & Statistical Analysis, and Functional Analysis. Fig.1b summarizes the workflow of the software. Also, Fig.1a indicates the homage of the app, including the corresponding tabs to analysis steps.

**Fig. 1.**
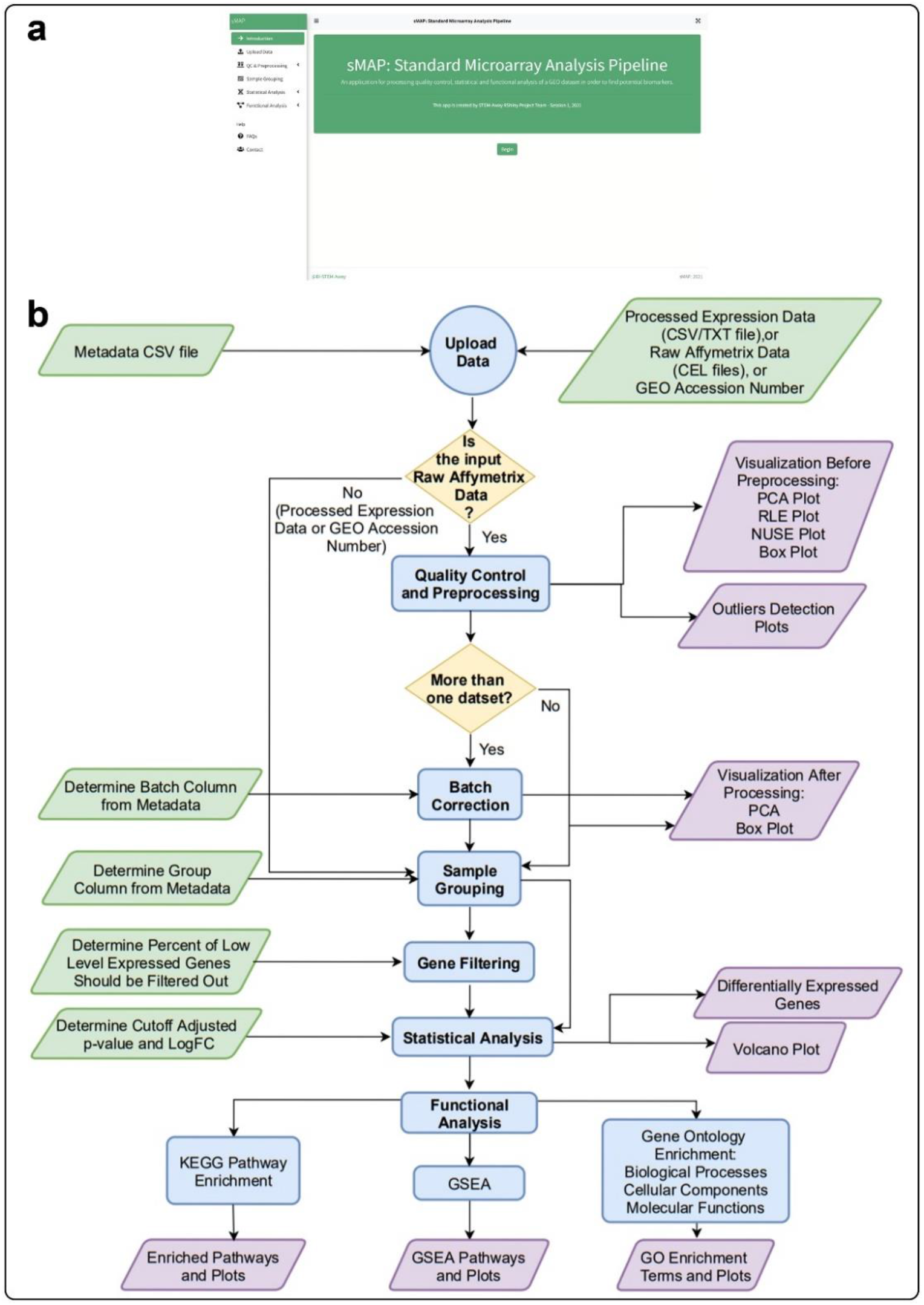
**(a)** Hobmepage of sMAP **(b)** Workflow of sMAP

### Software Implementation

sMAP application is supported by the R Shiny framework. The bs4Dash package is used for designing the user interface. The four major sections in sMAP, i.e. Data Importation, Quality Control & Preprocessing, Statistical Analysis, and Functional Analysis, are implemented using Bioconductor libraries and API functions. In addition to the sMAP web application which is currently hosted on an AWS cloud EC2 instance (t3.medium), users can also run an instance of sMAP on their local computer either in R or by running a docker container with sMAP’s docker image (https://hub.docker.com/r/samuelbharti/smap) to analyze large datasets. The latest hosting URLs of sMAP and a comprehensive tutorial can be found at https://github.com/BI-STEM-Away/sMAP.

### Data Importation

sMAP supports various data types and file formats as input and accordingly the user is instructed to perform quality control or to proceed to sample grouping. When starting with raw CEL files as the input, the user can perform quality control analysis steps suited for raw data. There are four supported data input types in sMAP; uploading an expression data file in .csv or .txt format, uploading raw Affymetrix data in CEL file format, entering a GEO accession number ID or selecting sMAP’s demo dataset. sMAP supports expression data from the Affymetrix Human Genome 1.0 ST Array or from the Affymetrix Human Genome U133 Plus 2.0 Array. For all input types other than the demo data, a metadata file must be uploaded in the csv file format with the first column containing the names or IDs of samples, and subsequent columns containing experimental features of interest or batch assignments if the data consists of various batches. For the demo dataset, the metadata and other inputs are preloaded.

### Quality Control & Preprocessing

When starting with a raw dataset (CEL files) as the input in sMAP, the user is able to perform all basic quality control (QC) analysis steps. QC analysis includes normalization, batch correction, and outlier identification and removal, complemented with different visualizations. The normalization of data can be performed using any of the three methods supported in sMAP: RMA, GCRMA, and MAS5. Following normalization, the user can perform batch correction to ensure the comparability of data from multiple dataset. The user can specify batch assignments for the samples by inputting which column of the metadata file contains the batch assignments. The data before and after normalization and batch correction are shown to users with supported visualization of NUSE (Normalized Unscaled Standard Errors), RLE (Relative Log Expression), PCA (Principal Component Analysis) plots, and boxplots of logarithm-scaled expression values. NUSE and RLE plots are generated using a probe-level model of the raw data. The last step in QC analysis is the detection of potential outliers for which sMAP allows the user to choose from three statistical methods: “KS (Kolmogorov-Smirnov) test”, “sum”, and “upperquartile’’ given by the ArrayQualityMetrics package. Detected outliers can then be removed by user selection and the final expression matrix and metadata are updated.

### Sample Grouping & Statistical Analysis

After importing data into sMAP and performing QC analysis (depending on the input data type), the users can perform sample grouping by selecting the column with grouping information from the metadata. The users can then optionally perform gene filtering before calculating DEGs using the limma package in R [13]. Genes are annotated based on the type of microarray used, and missing values will be removed [14]. sMAP also provides a parameter threshold of log fold change and adjusted p-value to filter the DEG result. The resulting table is used to generate a dynamic volcano plot with cutoff parameters and an option to toggle gene labels.

### Functional Analysis

sMAP also supports functional analysis methods using various libraries and API functions. The current implementation included GO (Gene Ontology) annotation, KEGG (Kyoto Encyclopedia of Genes and Genomes) enrichment analysis, and GSEA (Gene Set Enrichment Analysis). The results are displayed in dot plots, bar plots, and enriched pathway tables. Each output can be filtered using a p-value cut-off and is downloadable for use in publications.

## Case Study

To demonstrate sMAP functionalities, we analyzed the gene expression profiles of 17 cancerous and non-cancerous colorectal cancer paired samples from the dataset GSE32323 available in the GEO database. We compared the findings obtained from sMAP with the previously published study by Hu et al [15]. Hu et al. had analyzed the same samples from the GSE32323 dataset using a similar pipeline to the sMAP pipeline. The raw CEL files were imported (Fig. 2a) and a custom metadata file was created to import in sMAP. This data has also now been installed in sMAP as one of the demo datasets.

**Fig. 2.**
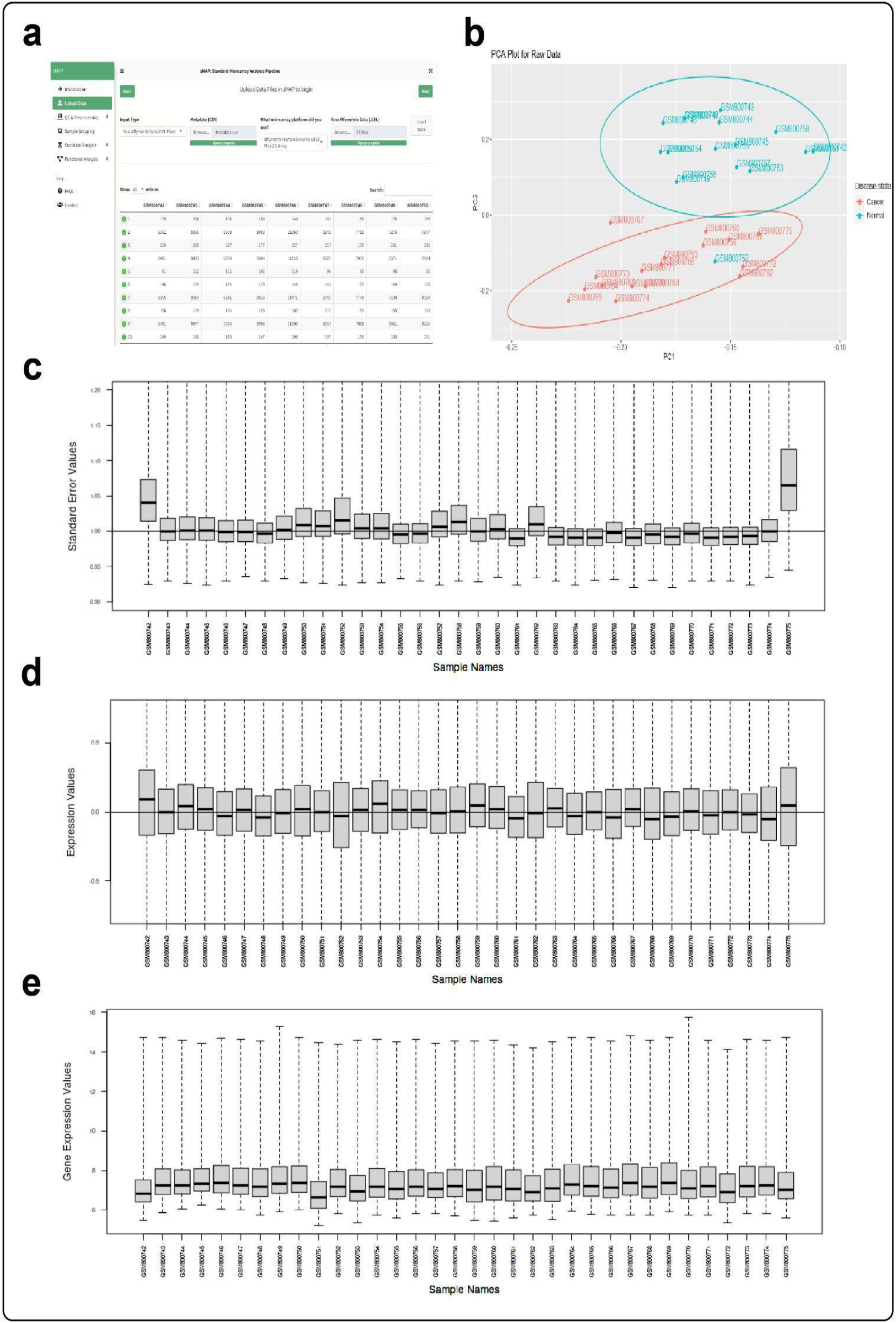
The analysis steps and plots of 17 paired samples from the GSE32323 dataset by the sMAP related to “Upload Data” tab and “Visualization Before Preprocessing” sub-tab: **(a)** Table of expression data after uploading CEL files **(b)** PCA of data before conducting preprocessing. **(c)** NUSE plot that effectively compares the probeset standard errors between samples. **(d)** RLE plot that shows the relative intensity values between samples. **(e)** Box plot that indicates the distribution of gene expression in samples.

sMAP detected two samples, GSM800775 and GSM800742, as outliers by the KS method (Fig. 3a and 3b). These outliers can also be recognized by looking at RLE and NUSE plots (Fig. 2c and 2d). However, Hu et al. had not removed outliers, so these outliers were not removed in our analysis to maintain the similarity between Hu et al.’s pipeline and the pipeline used in our case study. Also, PCA plot of data was constructed for further quality control (Fig. 2b). The RMA method was used for normalization and background correction, and the results were visualized using a boxplot (Fig. 3c) and a PCA plot (Fig. 3d). Samples were grouped by disease state using the Sample Grouping tab. An FDR (False Discovery Rate) correction method was used to correct p-values, and thresholds of adjusted p-value <0.05 and |Log Fold Change|>1 were applied on genes to obtain DEGs. These genes were visualized in the volcano plot (Fig. 3f). Hu et al. had performed KEGG pathway enrichment analysis and Biological Process GO term enrichment analysis. sMAP was used to perform these same analyses. We applied an adjusted p-value cutoff of p < 0.05, just as Hu et al. Fig. 3g and Fig. 3h indicate the results of these steps. Moreover, sMAP was used to perform GO enrichment analysis for Molecular Functions and Cellular Components with an adjusted p-value cutoff of p<0.05, and GSEA with the app’s default adjusted p-value threshold of p<0.05 (Fig. 4).

**Fig. 3.**
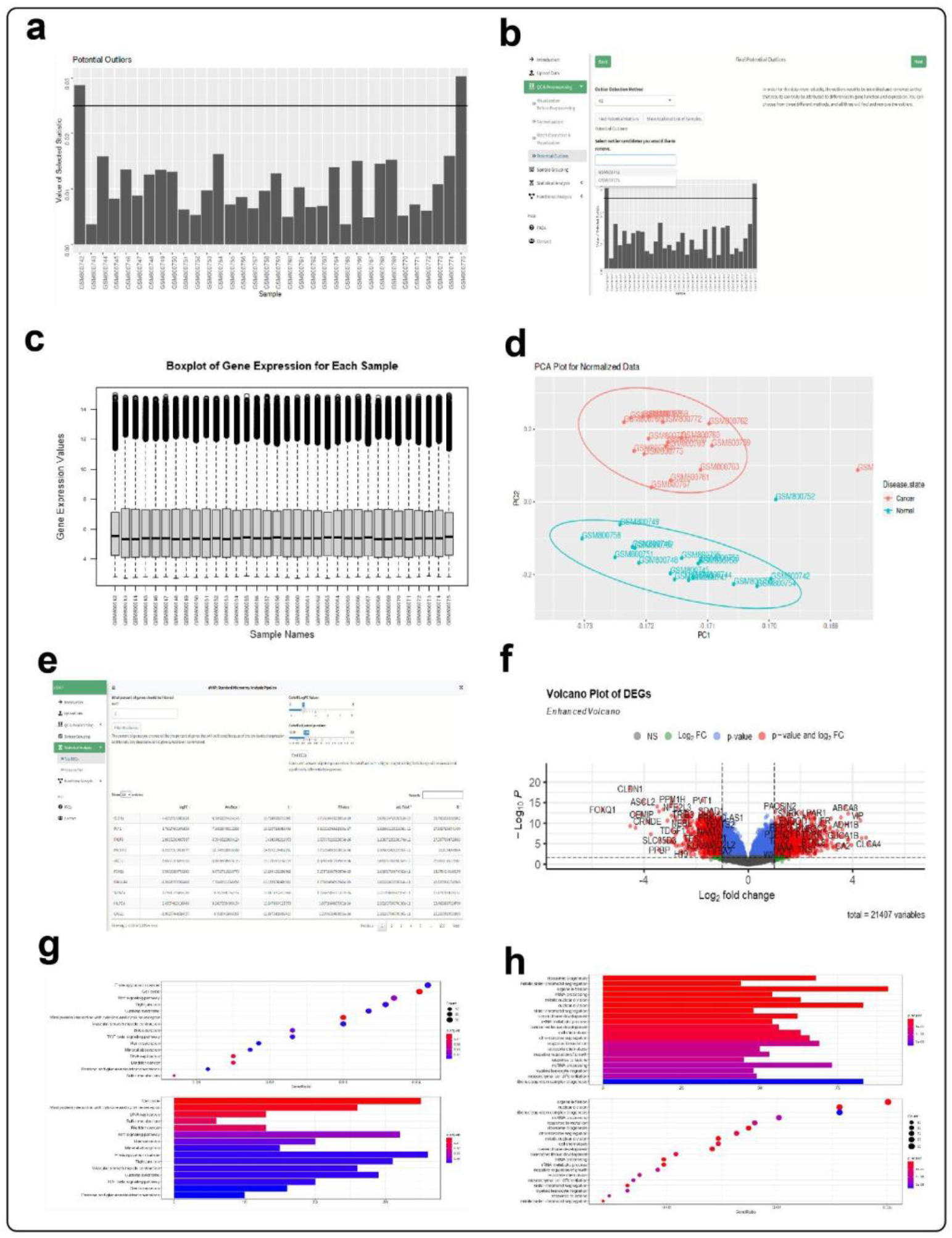
The analysis steps and plots of 17 paired samples from the GSE32323 dataset by the sMAP related to” Batch Correction & Visualization’’ and “Potential Outlier” sub-tabs and “Sample Grouping,” “Statistical Analysis,” and “Functional Analysis” tabs: **(a)** Results of applying KS outlier detection method on data (the samples that pass through the threshold in the bar plot are known as outliers). **(b)** The app shows the candidate outliers. The user can select them and click on the “Show Updated List of Samples’’ button to remove the outliers. **(c)** Boxplot after preprocessing by RMA method **(d)** PCA plot after preprocessing **(e)** List of (DEGs) after applying FDR-corrected p-value threshold of p<0.05 and |log fold change|>1 **(f)** Volcano Plot which is a scatter plot of the distribution of genes after differential expression analysis by log2 fold change and -log10 of the adjusted p-value. The vertical and horizontal lines on the plot respectively represent thresholds specified by the user. The green dots represent the genes that pass through the log fold change threshold, and the blue ones indicate the genes that pass through the adjusted p-value threshold. The red dots are the genes that meet both criteria (log fold change and adjusted p-value). **(g)** Enriched KEGG pathways are shown by the bar plot and the dot plot. The bar’s length and dot’s size indicate the counts of genes that enrich the corresponding KEGG pathway (count of core enrichment genes). In the dot plot, GeneRatio values are plotted on the x-axis, defined as the count of core enrichment genes divided by the count of pathway genes. **(h)** Enriched Gene Ontology terms (Biological Processes) are shown by the bar plot and the dot plot. The bar’s length and dot’s size indicate the counts of genes that enrich the corresponding Biological Process (count of core enrichment genes). In the dot plot, GeneRatio values are plotted on the x-axis, defined as the count of core enrichment genes divided by the count of Biological Process genes.

**Fig. 4.**
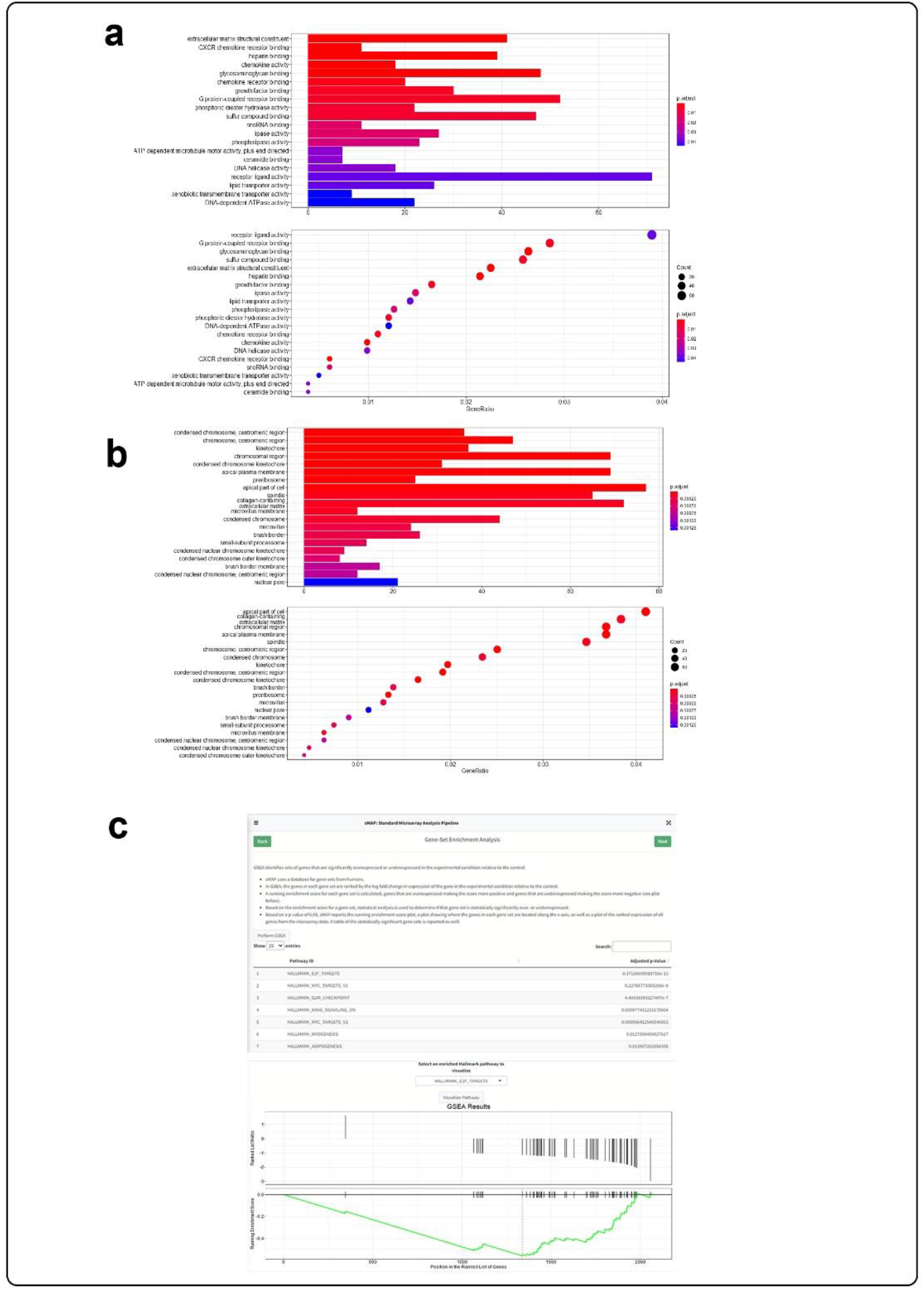
The analysis steps and plots of 17 paired samples from the GSE32323 dataset by sMAP related to “Gene Ontology Enrichment” and “Gene Set Enrichment” sub-tabs: **(a)** Enriched Gene Ontology terms (Molecular Functions) are shown by the bar plot and the dot plot. The bar’s length and dot’s size indicate the counts of genes that enrich the corresponding Molecular Functions (count of core enrichment genes). In the dot plot, GeneRatio values are plotted on the x-axis, defined as the count of core enrichment genes divided by the count of Molecular Functions genes. **(b)** Enriched Gene Ontology terms (Cellular Components) are shown by the bar plot and the dot plot. The bar’s length and dot’s size indicate the counts of genes that enrich the corresponding Cellular Components (count of core enrichment genes). In the dot plot, GeneRatio values are plotted on the x-axis, defined as the count of core enrichment genes divided by the count of Cellular Components genes. **(c)** GSEA uses HALLMARK gene sets, which represent well-defined biological processes or states, and looks to see whether they are enriched in the user’s experimental dataset compared to the control dataset. Those biological processes or states, along with their adjusted p-value, are indicated in a table in sMAP. The user can choose the number of biological processes or states to be shown in the plot and select one of them to be shown on the plot.

Hu et al. found 646 upregulated and 638 downregulated DEGs. The analysis by sMAP resulted in 985 upregulated and 1040 down-regulated DEGs (Supplementary File 2). When comparing the symbol names of the top 20 DEGs, when in ascending order based on adjusted p-value, in Hu et al.’s study and sMAP, 8 common DEGs were found: ASCL2, CLDN1, FOXQ1, AJUBA, ABCA8, MYC, HILPDA, and CHGA. Of the top 20 DEGs found in Hu et al.’s study, 18 of them were found to be DEGs in sMAP. Hu et al. also reported that the DEGs were enriched in 169 GO terms and 23 KEGG pathways. Among the 21 most significant KEGG pathways stated in Hu et al.’s article, 5 were also enriched in sMAP: “Cell cycle,” “DNA replication,” “Sulfur metabolism,” “Mineral absorption,” and “Wnt signaling pathway.” The comparison of the 10 most significant GO Biological Processes enriched terms reported by Hu et al. and the 15 most significant GO terms of sMAP resulted in 2 common GO terms: “Mitotic nuclear division” and “Negative regulation of growth”. The consistencies in results compared to a previously published study show the accuracy and reliability of sMAP’s results.

In order to further demonstrate sMAP’s abilities to identify significant pathological biomarkers and to integrate data from more than one dataset, another case study was performed with paired lung cancer samples from two different datasets (Supplementary File 1).

## Discussion and Conclusion

Microarray analysis is commonly used to extract functional genomics information from raw data. Early-stage researchers can benefit from microarray analysis but may not have the computational background needed to analyze their data thoroughly.

sMAP was developed as an interactive microarray analysis tool available for both online and offline use to aid early-stage researchers in the field of biology, bioinformatics, or similar. It can also provide new and significant insights from downstream analysis. Although we included most of the basic packages and methods required in a standard microarray analysis pipeline, we acknowledge there are more advanced methods and visualizations that can be implemented. For instance, supporting more input data formats, annotation packages, and enrichment analysis tools would all result in a more robust app. The large file sizes of raw microarray datasets may limit the use of sMAP’s web server, in which case the user can install/run sMAP on their local computer.

With an emphasis on interactivity and explanatory guidance, sMAP holds the potential to serve as an introductory tool to the world of microarray analysis, in which the user is able to derive a fundamental understanding of statistical and functional analysis of gene expression data. For early-stage researchers with limited computational proficiency, sMAP serves as a segue into more complex analytical processes. Thus, sMAP’s design considerations for early-stage researchers and public availability can be seen as a means of making research more accessible by increasing the scope of people who are able to draw biological conclusions from genomic data.

## Data Availability

### Documentation

sMAP users are provided with in-depth documentation in order to understand the customizable features and output results of the application (https://bi-stem-away.github.io/sMAP_doc/). Further details and links to sMAP’s GitHub repositories can be found at the project’s website (https://bi-stem-away.github.io/sMAP/).

**Table 1** sMAP supports four input types that either contain raw or preprocessed expression data.

### Supplemental Files

1. Supplementary File 1: Additional case study using sMAP.
2. Supplementary File 2: Table of Differentially Expressed Genes from both case studies.

## Author Contributions

AV and SR wrote the manuscript. NK and MM performed and wrote case studies, provided figures, performed literature validation, and helped in manuscript writing. SB designed, implemented, and deployed sMAP software and helped in manuscript writing. NK helped in the development of sMAP. All authors read, edited, and approved the manuscript.

## Acknowledgments

All authors thank and acknowledge STEM-Away Bioinformatics interns who shared ideas and discussions that made sMAP possible: Disha Chauhan, Roman Ramirez, Aditi Verma, Huikun Li, Shreya Vora, Ivan Lam, and Modupe Arotiba. We would also like to thank STEM-Away founder Dr. Debaleena Das for providing STEM-Away platform support to all Bioinformatics interns and AWS cloud resources to host sMAP web server.

