## Supplementary File 1 for "sMAP: An interactive microarray data analysis tool for early-stage researchers"

**Supplementary Case Study**

**Introduction**

To illustrate sMAP’s utility in identifying significant pathological biomarkers, samples were taken from two datasets. First, samples were taken from a data set of 120 paired samples of tumor and adjacent normal lung tissue specimens from nonsmoking females in Taiwan (GEO accession number GSE19804). Lu, et al., analyzed these samples to identify genes associated with the progression of lung cancer in nonsmokers and identified downregulation of *SEMA5A* in cancerous tissues, a gene associated with axonal guidance and a potential target for future therapeutics [1]. Next, 102 samples were taken from a dataset of 246 samples of lung tissue, 226 being lung adenocarcinomas and 20 samples being normal lung tissue (GEO accession number GSE31210). These 102 samples were obtained from non-smoking females and were analyzed together with the 120 samples from the GSE19804 dataset using sMAP. Okayama, et al., analyzed the expression profiles of the 226 samples and found that the samples could be clustered into *ALK*-positive and *ALK*-negative adenocarcinomas based on the presence of a mutated *ALK* gene, and that there was a relationship between the presence of the mutation and the severity of the individual’s prognosis [2]. Both datasets used the Affymetrix Human Genome U133 Plus 2.0 Array to obtain the expression data. Samples from each dataset were analyzed together to ensure that enriched genes and pathways significant to female non-smoking lung cancer could be identified with greater generalizability than what could be determined from analyzing a single dataset. This case study examines the results of analyzing the combined dataset.

**Importing Data and Quality Control & Preprocessing**

A total of 222 CEL files were inputted to sMAP along with the corresponding metadata file. The expression data after uploading the data is reported in a table on sMAP (Fig. 1a). The uploaded metadata file consists of a table with three columns, the first columns containing all of the sample accession numbers. The second column, titled “Disease State,” assigns each sample to a “Cancer” or “Normal” experimental group based on whether the sample was taken from a cancerous tissue or a normal, healthy tissue. The third column, titled “Batch,” assigns each sample to the dataset from which it was obtained. A value of 1 in this column indicates that the sample was taken from the GSE19804 dataset and a value of 2 in this column indicates that the sample was taken from the GSE 31210 dataset.

To begin quality control, the NUSE, RLE, PCA, and boxplot visualization options were used before normalization and batch correction (Fig. 1b-1e). From the RLE and NUSE plots, it seems that there are multiple samples with RLE and NUSE distributions that are different than the other samples. The NUSE plot highlights the presence of batch effects as the samples from the GSE19804 dataset, which are located on the left side of the plot, have higher median values and greater spread for NUSE values than the samples from the GSE31210 dataset. The boxplot visualization of the raw samples shows great variability between samples within the GSE19804 dataset and between samples of the GSE19804 dataset as compared to samples of the GSE31210 dataset. These visualizations thus show that for this data, batch effects are causing notable variability in expression values and may affect statistical analysis if not accounted for.

To process the data, the RMA normalization method was used for the combined dataset as this method is commonly used and proved to be effective in getting more similarly distributed samples, as indicated by the visualizations after normalization and batch correction (Fig. 2a and 2b). Batch correction was performed to correct for differences due to the fact that samples were obtained from different datasets, including different instruments and experimenters. The boxplot shows that the variation between batches has been reduced (Fig. 2a). The PCA plot indicates that the treatment groups are still not completely separable after pre-processing, which could indicate that there are limitations in using the GSE19804 and GSE31210 datasets together (Fig. 2b). sMAP is a useful tool for discovering new biomarkers that can then be further studied and verified, thus making this limitation acceptable for the sake of identifying potentially meaningful genes and pathways.

To determine quantitatively if any outliers are present in the combined data set, a plot comparing the KS statistic was computed and visualized for each sample (Fig. 2c). In the KS plot, samples GSM494574, GSM494590, GSM494591, GSM494594, GSM494596, GSM494601, GSM494649, GSM494650, GSM494654, GSM494656, GSM494657, GSM494660, GSM494661 were all determined to be outliers. These samples were removed from the dataset before continuing to statistical analysis.

**Obtaining Differentially Expressed Genes (DEGs)**

In the metadata file, each sample was assigned to a group under a column called “Disease State,” cancerous samples being a part of the group “Cancer,” and healthy samples being a part of the group “Normal.” This grouping was used to perform statistical analysis of differentially expressed genes and was specified using the “Sample Grouping” tab of sMAP. After grouping the samples, the DEGs were obtained by using the sMAP’s default cutoff values, the log fold change cutoff value being 1 and the adjusted p-value cutoff value being 0.05. The default values were selected as previous work using each gene set used different statistical tests and cutoff values than each other and sMAP. Genes expressed at low levels were not filtered for this dataset as it was desirable to consider all potentially significant biomarkers. sMAP found 619 DEGs, which were visualized in a volcano plot (Fig. 2d).

sMAP found *SEMA5A* to be downregulated, as Lu, et al. found, and also found *SEMA6A*, *SEMA6G,* and *SEMA3G* to be significantly downregulated, offering additional support for Lu, et al.’s finding that semaphorin gene expression is altered in lung cancer patients. Moreover, sMAP identified the top ten upregulated and top ten downregulated genes from a study conducted by Lv, et al. comparing healthy and cancerous samples from only the GSE19804 dataset as being upregulated and downregulated, including *COL10A1* and *HS6ST2*. In addition to reproducing results from previous work, sMAP’s ability to correct for batch effects and be applied to multiple datasets allowed for the identification of genes that may be associated with NSCLC but have neither been reported in work analyzing the GSE19804 dataset nor in work analyzing the GSE31210 dataset. For example, the gene *GCNT3*, which is associated with multiple cancer types including NSCLC, was identified by sMAP as one of the most upregulated genes in the combined dataset. [3].

**Functional Analysis**

KEGG Pathway Enrichment Analysis revealed cellular pathways enriched in the cancerous tissue samples (Fig. 2e). The most enriched pathways identified by sMAP include pathways associated with cytokine signaling and cell adhesion molecules. The malaria KEGG pathway was also found to be significantly enriched, and previous work has shown that lung cancer tumor growth can be limited by infection by a malaria parasite, suggesting that exploring the connection between genes associated with malaria and lung cancer could be a promising therapeutic strategy [4].

The GO Enrichment Analysis seemed to yield similar results to those obtained from conducting GO Enrichment Analysis on the Lu, et al. data set. However, one of the terms under the category “cellular components” was found to be enriched in the combined data set but not when only the Lu, et al. data set is used. The “membrane raft” term was enriched for the combined data set (Fig. 2f-2h). Lipid rafts in the cell membrane are usually specialized domains in the membrane with a particular function, and in the case of lung cancer, domains are thought to play a role in the formation of exosomes that are part of cellular communication and protein transport, and are thought to be important biomarkers for many types of tumors, including those associated with lung cancer [5]. Findings from sMAP seem to confirm the important role exosomes may play in tumorigenesis, suggesting that genes associated with lipid raft formation, particularly those that form exosomes, should be further studied.

The Gene Set Enrichment Analysis (GSEA) of the data shows that gene sets associated with rapid cell growth, including the G2M checkpoint gene set, the E2F gene set, and the mitotic spindle gene set, are upregulated (Fig. 3a-3e). The epithelial to mesenchymal gene set is also enriched, and the process of epithelial to mesenchymal transition of tumor cells is associated with metastasis. The enriched gene sets thus are consistent with previous analysis of cancer cell behavior.

**Conclusion**

sMAP’s ability to correct for batch effects allows for multiple datasets to be analyzed together, allowing the user to potentially identify novel biomarkers or genetic pathways of interest for a given application. sMAP was used to analyze 222 samples taken from two datasets of females with lung cancer, GSE19804 and GSE31210. Analysis in sMAP resulted in 619 DEGs, many of which were consistent with previous work analyzing the GSE19804 dataset as well as some DEGs not previously reported. Functional analysis identified KEGG pathways, gene ontology terms, and hallmark gene sets that were enriched in the datasets and that were consistent with previous work. sMAP is thus a promising tool for both educating users about transcriptomic analysis as well as for analyzing microarray data to gain new insights about disease.

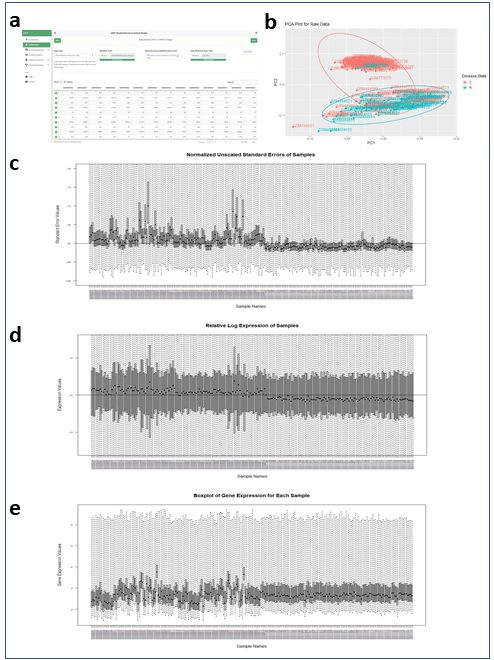

##### **Fig. 1** The raw CEL files for the selected samples were imported and processed in sMAP. sMAP enables the user to see the need for batch correction for this data. **(a)** 222 CEL files and a metadata file were input to sMAP. **(b)** A PCA plot of the raw data showed that the samples separate into cancerous (denoted by “C” in the legend) and normal (denoted by “N” in the legend) groups. **(c)** The NUSE plot revealed variation between samples based on batch, the samples from GSE19804 having higher values than the samples from GSE31210. **(d)** The RLE plot showed differences in the samples’ range of expression values, samples from GSE19804 having slightly larger ranges than samples from GSE31210. **(e)** The boxplot of log scaled expression values revealed variation between samples within the GSE19804 batch as well as between samples from the GSE19804 batch compared to samples from the GSE31210 batch.

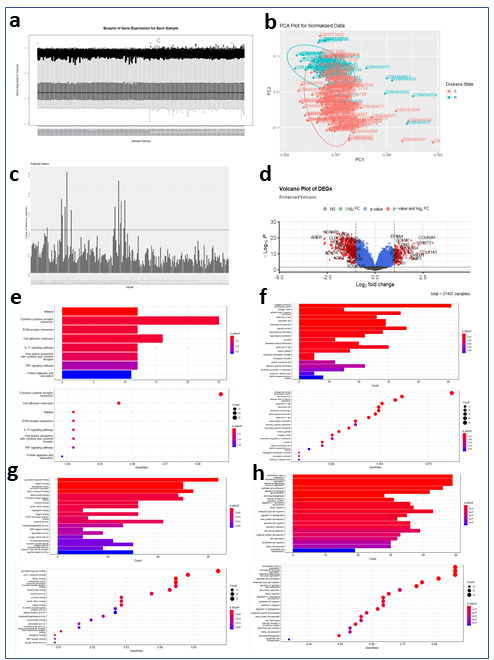

##### **Fig. 2** After RMA normalization and batch correction, the data was visualized and outliers were identified. The processed data was visualized using: **(a)** a boxplot, showing that the data is normalized and that there is less intra- and inter-batch variation, and **(b)** a PCA plot, showing some separation between cancerous and normal samples. **(c)** Outliers with a KS statistic past the threshold were removed. **(d)** A volcano plot was used to visualize the 619 DEGs. **(e)** KEGG analysis revealed 8 enriched pathways, including cytokine signaling and cell adhesion. **(f)** GO analysis of Cellular Components resulted in terms involving the cell membrane to be enriched, among others. **(g)** GO analysis of Molecular Functions resulted in terms involving molecular binding to be enriched, among others. **(h)** GO analysis of Biological Processes resulted in terms involving extracellular structure and cell proliferation to be enriched, among others.

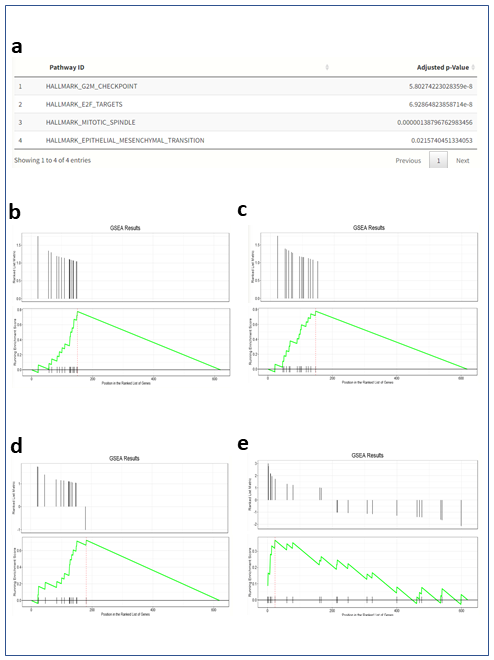

##### **Fig. 3** GSEA was conducted. **(a)** Four Hallmark gene sets were found to be enriched: **(b)** G2M Checkpoint, **(c)** E2F Targets, **(d)** Mitotic Spindle, and **(e)** Epithelial to Mesenchymal Transition.
